# Intelligible distracting speech disrupts early auditory attention

**DOI:** 10.64898/2026.08.19.745879

**Authors:** Benjamin N. Richardson, Maanasa Guru Adimurthy, Christopher A. Brown, Antje Ihlefeld, Merri J. Rosen, Barbara G. Shinn-Cunningham

## Abstract

Intelligible speech disrupts selective auditory attention more than an unintelligible stream. However, low-level acoustic features of intelligible speech are relatively similar to target speech, confounding results. While controlling acoustic similarity and limiting energetic masking, we examined how masker intelligibility affects behavior and electroencephalography (EEG). Normal hearing listeners detected color words within a target stream of randomly timed words while ignoring an ongoing masker. Maskers were either spoken by the same or a different talker and comprised either isochronous sequences of intelligible words or temporally scrambled versions. Scrambled maskers either lacked broadband energy changes over time (Experiment 1) or were amplitude modulated to have the same energy profiles as intelligible, isochronous maskers (Experiment 2). In both experiments, scrambled maskers yielded better performance than intelligible maskers. For intelligible maskers, performance was better for different compared to identical talkers. EEG responses paralleled behavior: target-evoked onset responses were larger for scrambled than for intelligible maskers, particularly for identical talkers. Later target recognition responses were larger for color than other target words but unaffected by masker type or talker. Even when low-level acoustic features were carefully matched, intelligible maskers impaired auditory attention and reduced target-evoked neural responses more than scrambled maskers, implicating early sensory filtering.

## I. Introduction

Intelligible speech maskers are known to impair speech perception more strongly than maskers that are acoustically similar but unintelligible (Kidd and Colburn, 2017). The increased perceptual interference from intelligible (compared to unintelligible) maskers is seen across a wide range of paradigms, including comparisons between speech and noise maskers (Best et al., 2020; Brungart and Simpson, 2002), manipulations of the number of talkers in multitalker babble (Brungart et al., 2001; Kurthen et al., 2021; Van Engen et al., 2014), differences in language familiarity (Van Engen and Bradlow, 2007), spectral or temporal filtering (Pollack, 1952, 2005; Rennies et al., 2019), noise vocoding (Scott et al., 2006; Strelnikov et al., 2011), and time reversal or scrambling of speech (Summers and Roberts, 2020). These results highlight that a masker comprising structured, intelligible speech interferes with perception of target speech more than many other types of maskers.

Despite this body of evidence suggesting that intelligibility influences how strongly a masker interferes with perception of a target speech stream, most past studies fail to disentangle contributions of masker intelligibility from acoustic similarity of the masker and the target speech. For example, increasing the number of talkers in babble changes the density of energy within individual time-frequency regions; filtering or vocoding alters spectrotemporal detail in the signals; and time reversal modifies temporal envelopes. This leaves open whether previously observed effects of intelligibility truly reflect linguistic interference or differences in acoustic similarity between target and masker.

One specific form of target-masker similarity known to influence perceptual interference from a masker is similarity in talker identity. Competing sounds that share vocal characteristics such as fundamental frequency, vocal tract length, or timbre are more difficult to segregate and more likely to interfere with target perception (Best et al., 2007; Brungart et al., 2001; Darwin, 2008; Darwin and Carlyon, 1995; Zekveld et al., 2014). When target and masker are spoken by the same talker, stream segregation is weakened, making listeners more likely to confuse target and masker content and report content of the masker as belonging to the target stream (Shinn-Cunningham et al., 2017; Shinn-Cunningham, 2008). These effects demonstrate that intelligibility of a target stream depends on whether the auditory system can successfully segregate the target and masker streams.

While behavioral outcomes have been instrumental in characterizing masking by intelligible (and unintelligible) speech, neural measures can reveal whether this interference arises at early sensory stages or later stages of selection and recognition. Animal electrophysiology consistently shows that attention modulates sensory responses in the auditory cortex. Attention sharpens sensory gain and selectivity in primary and other regions of auditory cortex (Hockley and Malmierca, 2024; McGinley et al., 2015; Niwa et al., 2012, 2015; Winkowski et al., 2018). Invasive neural recordings show that higher order human auditory cortex encodes basic acoustic features of a stimulus (Pasley et al., 2012). Importantly, cortex also preferentially represents features of attended sounds in favor of unattended ones (Mesgarani and Chang, 2012; Zion Golumbic et al., 2013).

Comparable effects of gain and selectivity tuning have also been shown through noninvasive magnetoencephalography (de Vries et al., 2021; Woldorff et al., 1998), functional magnetic resonance imaging (Alho et al., 2014; Lee et al., 2013, 2014; Woldorff et al., 1993), and electroencephalography (Kerlin et al., 2010; Power et al., 2012). EEG lacks the spatial precision of invasive recordings, but it noninvasively samples activity across broad, distributed regions, and its accessibility makes it well suited to the present study. Across imaging modalities and tasks, the convergent pattern suggests that attentional gain is not tied to any single privileged acoustic cue, but instead flexibly exploits whatever feature reliably distinguishes the target from competing sound sources (talker identity, spatial location, etc.). This is also true of models: a network trained with stimulus-computable feature gains reproduced human-like selective listening without being trained to mimic human behavior (Griffith et al., 2026). Taken together, these findings indicate that selective attention operates in part by biasing early sensory encoding in favor of target sounds rather than solely resolving competition at later decision stages.

To dissociate how gain control in auditory cortex drives intelligibility by modulating early sensory competition compared to later stage cortical processing, we analyzed event-related potentials (ERPs) recorded using electroencephalography (EEG). A particularly useful window into attentional processes, the P1-N1 complex of the auditory ERP is sensitive to attentional modulation and reflects gain control in auditory cortex, with attended sounds typically evoking larger response magnitudes than unattended sounds (Choi et al., 2014; Hillyard et al., 1973; Noyce et al., 2021). Later latency ERP components like the P300 reflect higher order cortical processing. The P300 reflects detection or categorization of task-relevant, attended sound events and is associated with later stages of stimulus evaluation and decision-making (Picton, 1992; Polich, 2007). P300 amplitudes are sensitive to the probability of a target stimulus, task demands, and whether a participant is attending to the stimulus (Polich, 1986). When observed together, the P1-N1 and P300 components allow dissociation between failures of early sensory filtering and failures of recognition or categorization.

Across two experiments in normal hearing listeners, we sought to determine how masker intelligibility and talker similarity influence selective auditory attention to a sporadic target stream of color and object words embedded within an ongoing, isochronous masker stream. We concurrently measured detection of target color words and evoked response components P1-N1 and P300. We contrasted different types of maskers to isolate the contributions of masker intelligibility and talker similarity while minimizing low-level acoustic confounds. Specifically, we presented target speech with ongoing masker streams that were either intelligible word streams, or temporally scrambled versions of these streams that preserved the distribution of short-term spectral content of the intelligible speech. We further contrasted maskers from the same talker as the target, or from a different talker. In a second experiment, we introduced an additional control on the temporal envelope of the scrambled-speech maskers.

We hypothesized that intelligible speech maskers would disrupt selective auditory attention more strongly than unintelligible, scrambled-speech maskers, even when the instantaneous spectral structure of the maskers (and thus the energetic overlap between target and masker) were carefully matched. We further hypothesized that this disruption would be exacerbated when the target and masker were spoken by the same talker, reflecting reduced effectiveness of attentional selection due to increased target-masker similarity. At the neural level, we predicted that early sensory responses to target onsets, indexed by the P1-N1 complex, would mirror behavioral outcomes. We expected that the P1-N1 elicited by a target sound would be smaller when the masker was difficult to ignore, that is, intelligible and spoken by the same talker. We also expected that the P300, as an index of target recognition, might also be affected by properties of the masker; we expected an intelligible, same-talker masker to produce a smaller P300 due to increased interference with target identification and selection.

We found that masker intelligibility and talker differences substantially modulated behavioral performance and reduced early sensory (P1-N1) responses to target sounds, consistent with a failure of effective sensory filtering with an intelligible or same-talker masker. In contrast, P300 responses associated with successful target color word recognition were largely insensitive to masker intelligibility and talker differences. These findings suggest that intelligible competing speech primarily interferes with selective attention at early stages of sensory-perceptual processing, limiting the likelihood that target sounds “get through” the attentional filter to reach recognition. By combining rigorous acoustic control with behavioral and neural measures, the present study helps clarify how intelligibility and talker identity shape the perceptual organization of competing speech and the neural mechanisms that support selective auditory attention.

## II. Materials and Methods

### A. Participants

Twenty normal hearing, native English speaking listeners participated in each of Experiments 1 (13 female, 4 male, 3 not reported, age 26.9 ± 1.53 SD) and 2 (10 female, 10 male, 0 not reported, age 20.4 ± 4.40 SD). This sample size is consistent with prior ERP studies of auditory selective attention in our lab and others that have reported reliable effects in early (P1–N1) and later (P300) components with similar group sizes. Normal hearing was confirmed with pure-tone audiometry: all participants had thresholds of 20 dB HL or better at octave frequencies from 250 Hz to 8 kHz in both ears. All listeners gave written informed consent prior to participation. All testing was administered according to the guidelines of the Institutional Review Board of Carnegie Mellon University.

### B. Color Word Detection Task

In both experiments, participants were instructed to monitor a target sound stream for color words and simultaneously ignore a masker stream (see Fig. 1). Participants were instructed to attend to the sporadic, randomly timed target stream (first row of Fig. 1) and ignore the ongoing, isochronous masker stream. Each 12-second block of target and masker streams comprised color or object word tokens. In both experiments, the masker stream was either words (second row of Fig. 1, Words masker) or temporally scrambled words (third and fourth rows of Fig. 1, Scrambled masker). The target and masker streams were either spoken by the same male talker, or different talkers.

**Figure 1.**
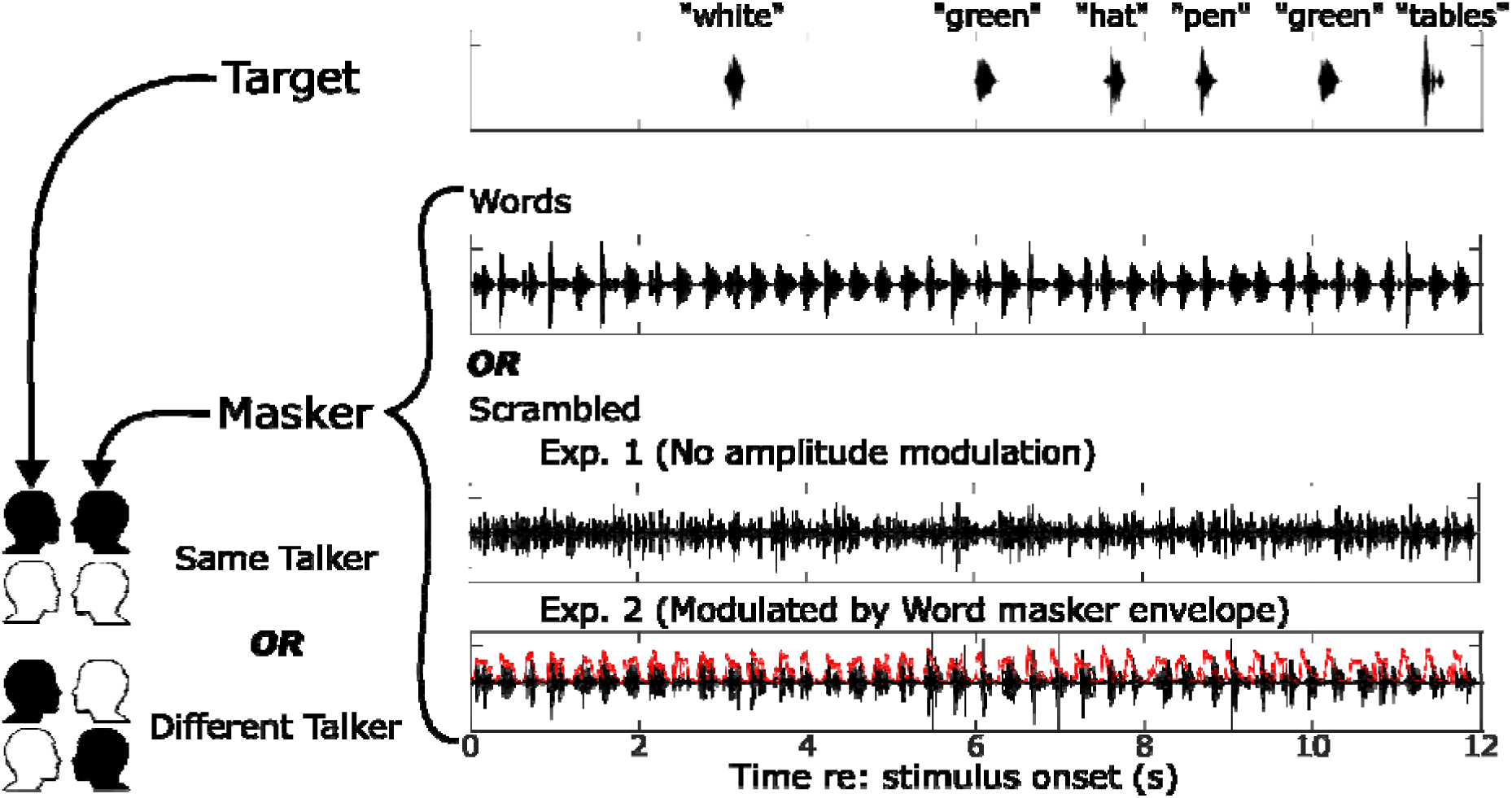
Diagram illustrating the target and masker streams for one block. Each 12-second block presented an ongoing masker stream (bottom rows) and a target sequence consisting of randomly timed target words. Participants were tasked with pressing a key whenever they heard a color word in the target. The masker stream within each experiment was either an isochronous sequence of words or a temporally scrambled version of the isochronous stream. In Experiment 1, the scrambling eliminated clear onsets and offsets. In Experiment 2, the scrambled masker was multiplied by an envelope derived from a word masker stream to reimpose rhythmic structure. In both experiments, the masker could either be from the same male talker or a different male talker as the target stream.

For each experiment, this yielded a 2 x 2 design, with independent variables of masker (Words, Scrambled) and talker (Same, Different). Thirty-six blocks of each condition were presented in each experiment, resulting in 144 blocks per experiment. The block presentation order was pseudorandom for each listener, logically organized into 36 “chunks.” Within each chunk, each of the four conditions was presented once, but within chunks, the order of conditions was randomized independently for each chunk and each listener.

Listeners were instructed to press a button when they detected a color word in the target stream (during each presentation block). The participant began each block by pressing a key, and sound began 1 second later. A fixation cross appeared on screen at the start of each block, on which participants were asked to maintain fixation.

### C. Stimuli and Procedure

Participants sat in a double-walled sound attenuating booth and listened to stimuli over Etymotic ER2 insert earphones (Etymotic Research, Elk Grove, Illinois). All stimuli were created in MATLAB version R2022b (The MathWorks Inc., Natick, Massachusetts) using custom written scripts and the AuditoryToolbox package version 2 (Slaney, n.d.). Both target and masker sound streams consisted of common English words, 300-ms in duration, spoken by one of two male talkers (Kidd et al., 2008). Word tokens were either colors {blue, green, red, white} or objects {bag, desk, glove, pen, table, toy, hat, card, chair, shoe, sock, spoon}. All stimuli were presented at 65 dB SPL.

The target stream comprised color and object words spoken by one of two male talkers. Between 3-5 color words and 3-5 object words occurred in the target stream, resulting in a total of 6, 8 or 10 words. The number of color words in a given block was equal to the number of object words. The order of the words in the target stream was chosen randomly, with replacement, except with the constraint that the same word could not occur simultaneously in the target and masker streams. The onset times of words in the target stream were also distributed randomly, except with the constraints that target words were at least 800 ms apart from each other, a target word onset did not occur within ±80 ms of the onset of a masker word. Target words were also restricted from occurring within the first second of the stimulus, or after the onset of the last masker word. With these constraints, the gap between target words ranged from 800 ms - 3.6 s, with an average of 1.4 s.

The masker stream was either a sequence of unscrambled words or a scrambled version of the word masker. As described below, scrambled masker streams were derived from masker word streams by temporally scrambling word streams. This procedure created scrambled maskers that were not intelligible, but that not only had the same long-term spectral content as a word masker stream, but also had the same distribution of short-term spectrotemporal structure (e.g., harmonic structure, formant structure, etc).

The word masker comprised 40 consecutive word tokens in an isochronous stream, resulting in a rhythmic sequence where word onsets appeared every 300 ms within a block; each block had a duration of 12 seconds (see second row of Fig. 1). The words in the masker stream were chosen randomly and independently for each block, with the additional constraints that no two consecutive words could be color words. Further, neither the first nor last words of the masker stream could be color words.

Each scrambled masker was created by first constructing an unscrambled masker, then temporally scrambling it. The unscrambled masker was divided into 25 ms windows. Each 25 ms time segment was then Hanning-windowed and overlapped by 50%. Then, windows were randomly shuffled in time over a 250 ms range (±125 ms), and overlap-added to create the scrambled signal. Scrambling at this time scale creates a masker that has a roughly constant energy envelope, removing syllabic structure while preserving the distribution of instantaneous spectra. Specifically, scrambling with a 25 ms window duration and a 250 ms shuffle radius affected temporal modulation rates between approximately 4 and 40 Hz, disrupting the syllabic-and phonemic-rate envelope structure and across-time statistical correlations. Because each window contained an unaltered snippet of the original waveform, however, F0 and voicing characteristics that fall at higher frequencies were largely preserved; only their position in time changed.

In Experiment 1, the output of the scrambling procedure served as the scrambled masker. Experiment 2 added one additional processing step to reinstate the rhythmic structure of the word masker stream, with an onset every 300 ms. To do this, we extracted the amplitude envelope of the unscrambled version of the masker using a 4th order low-pass Butterworth filter with a cutoff of 125 Hz and multiplied the scrambled masker with this envelope. Following this, the word-modulated scrambled masker was equalized so that it had the same root-mean-square intensity as its unmodulated version.

We were interested in the effects of intelligibility (scrambled vs. unscrambled maskers) and talker identity (same vs. different talker for target and masker) on performance. We therefore limited the influence of energetic masking by filtering target and masker streams into separate interleaved, non-overlapping frequency bands (Panel A, Figure 2). Each individual target and masker word was filtered into 16 adjacent frequency bands (9th order zero-phase Butterworth bandpass filters), logarithmically spaced between 300 Hz and 10 kHz. Then, target word tokens were reconstructed as the sum of the odd numbered frequency bands, while the masker stream was reconstructed as the sum of the filtered signals in even numbered frequency bands. All filtered words were then equalized in broadband root-mean-square energy before concatenating them to the stream, resulting in a 0 dB SNR.

**Figure 2.**
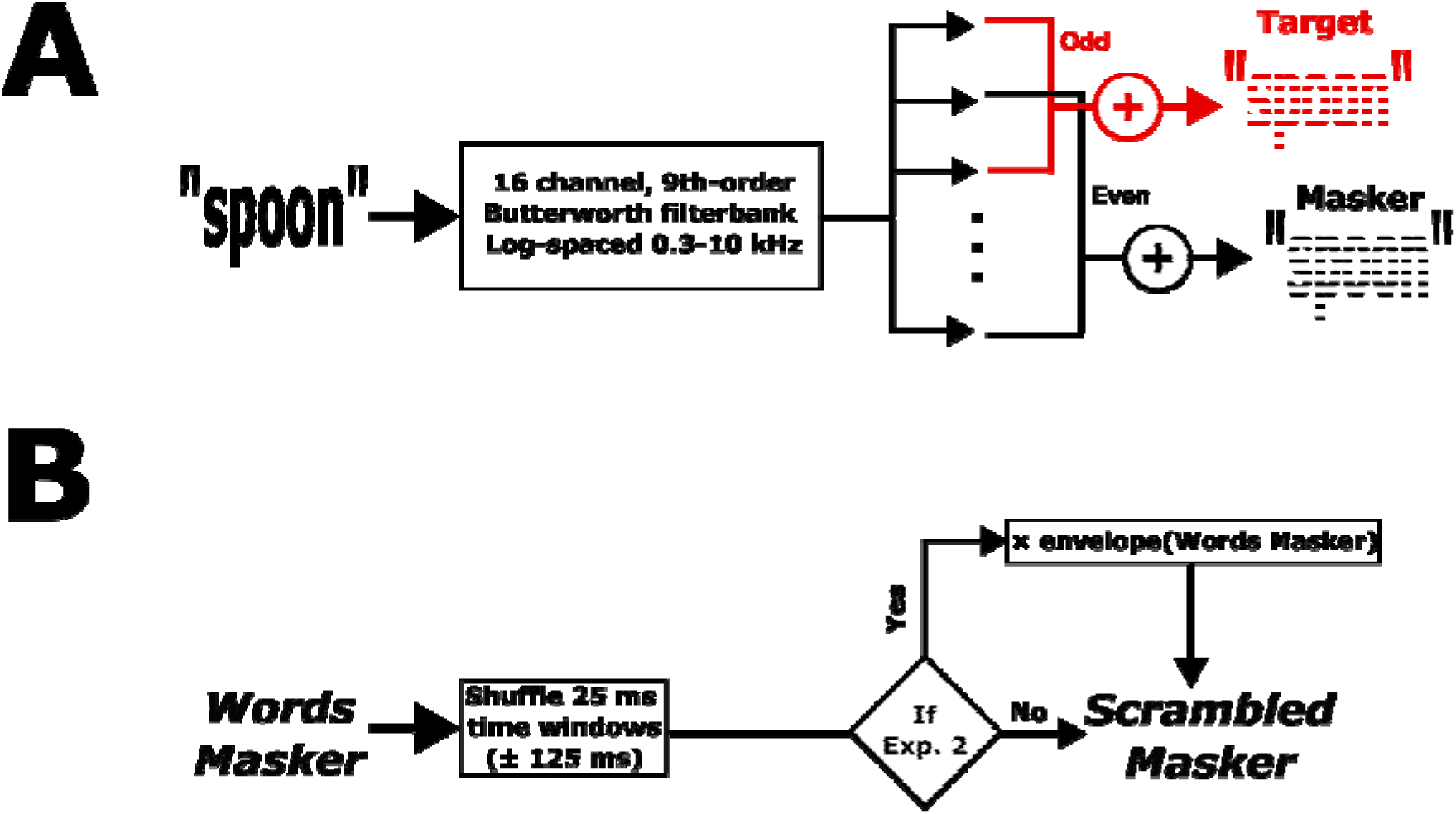
Controls for energetic masking and spectrotemporal properties. (A) To ensure minimal energetic masking between target and masker streams, each word token was filtered into 16 non-overlapping adjacent frequency bands. If that token belonged to the target stream, the odd numbered filterbank outputs were used to reconstruct the word token. If the word belonged to the masker, the even numbered outputs were used. (B) To ensure similar spectrotemporal properties of a Word masker and two types of Scrambled masker, the “scrambling” function in Auditory Toolbox (Slaney, n.d.) was used. Individual masker stream time windows were shuffled. For Experiment 2 only, the resulting masker was amplitude modulated using the envelope of its Words Masker counterpart.

### D. Behavioral Performance Analysis

We recorded the time of each button press with respect to the onset of each target word token to determine the rate of “hits” (correct responses to color words in the target stream), “within-stream false alarms” (incorrect responses to object words in the target stream), and “between-stream false alarms” (incorrect responses to color words in the masker stream, when the masker was words). Between-stream false alarms were only calculated when the masker was Words, because color words were not intelligible when the masker was Scrambled. We defined potential hit, within-stream false alarm, and between-stream false alarm windows as being from 300-1100 ms following word onset. Because target onsets occurred at least 800 ms apart, overlapping hit and within-stream false alarm response windows did not occur. Overlapping hit windows also did not occur because two color words could not occur consecutively in the target stream. However, in the case of conflicting potential response windows for between-stream false alarms, both potential windows were removed. Hit, within-stream false alarm, and between-stream false alarm rates were calculated as the total sum of key presses in each of the potential windows, divided by the total number of those windows. Rates were computed separately for each participant and each condition.

### E. EEG Data Acquisition and Analysis

A 32-channel Biosemi electroencephalography (EEG) system (“Biosemi EEG ECG EMG BSPM NEURO amplifier electrodes,” n.d.) recorded data throughout the task. Electrodes were arranged using the standard 10-20 layout. Data were acquired using BIOSEMI Actiview at a 2048 Hz sampling rate. Triggers were sent using a Cortech Triggy system (Cortech Solution, Wilmington, NC).

EEG data analysis was performed in MATLAB R2022b (The MathWorks Inc., Natick, Massachusetts) using the EEGLAB analysis package (Delorme and Makeig, 2004). First, EEG data were referenced to the mean of mastoid electrodes. Data were bandpass filtered between 1 and 30 Hz using a 2nd order Butterworth filter, and downsampled to 256 Hz. Next, the data were visually inspected to reject moments in time with excessive noise. Individual channel spectra were inspected and erratic channels were interpolated (Delorme & Makeig, 2004). Visual inspection of the components found using independent components analysis allowed us to isolate eye blinks, saccades, and motion artifacts. Signals were reconstructed after projecting out the relevant noise components. Across both experiments, an average of 1.1 ± 0.94 S.D. channels were removed and an average of 3.2 ± 1.2 SD components were projected out of the data. Data were epoched between 1 second prior to stimulus onset and 13 seconds after stimulus onset.

### F. Event-related potential (ERP) calculations

We isolated ERPs in response to words in the target stream and in the two masker streams with temporal structure (the word masker stream in both experiments and the word-modulated scrambled masker in Experiment 2). Individual ERP responses with deflections greater than ± 100 µV were rejected from further analysis (mean: 7.7 ERPs ± 3.7 S.D.). For each participant and condition, we computed the average ERP across remaining epochs in response to each color and object word onset in the target stream. Separately, we calculated the average response at every 300 ms interval during the stimulus, when a word onset occurred in the masker stream. All ERPs were baselined by subtracting the mean voltage during the 50 ms before these onsets. For ERPs for words in the target stream, only ERPs corresponding to hits and correct rejections were included in further analysis (see supplementary figure B).

We were interested specifically in whether target-word evoked ERPs reflected differences across conditions and for color vs. object words. Specifically, we examined the P1 to N1 depth, which represents the magnitude of auditory cortical sensory responses and is known to vary with attentional focus (Alho et al., 1994; Choi et al., 2014). We also calculated the P300 magnitude, which is larger in response to words that are task relevant (e.g., color target words compared to non-color target words) (Picton, 1992). The P1-N1 was calculated in a cluster of 8 frontocentral electrodes where auditory-evoked responses are maximal: Fz, FC1, FC2, C3, Cz, C4, CP1, and CP2. The P300 was calculated in a cluster of 8 parietooccipital electrodes, where this component is strongest: P3, Pz, PO3, O1, Oz, O2, PO4, and P4 (Başar-Eroglu et al., 2001). Each individual participant’s P1 and N1 peak times were first calculated by averaging the ERP to all target word tokens across all 144 blocks. Participant-specific P1 times were calculated as the time of the largest local maxima between 50 and 150 ms following word onset in the mean of frontocentral electrodes and N1 time as the time of the largest (most negative) local minima between 150 and 250 ms following word onset time, also in the frontocentral cluster. Participant-specific P300 times were found as the time of the largest local maximum between 250 and 600 ms in the parietooccipital cluster. We then analyzed how the P1-to-N1 and P300 response magnitudes at these participant-specific times varied with target word type, masker type, and talker identity. Specifically, each component’s magnitude was computed as the mean in either the frontocentral (P1, N1) or parietooccipital (P300) electrode cluster in a ± 20 ms time window around each participant’s times. To calculate the P1-N1 depth, we subtracted the P1 magnitude from the N1 magnitude for each participant, word type, and condition.

### G. Statistical analyses

For each measure of interest (hit rate, within-stream false alarm rate, between-stream false alarm rates, P1-N1 magnitude, and P300 magnitude), we used linear mixed effects models to test experimental effects. We fit separate models for Experiment 1 and Experiment 2. Behavioral models included fixed effects of Masker type (word, scrambled) and Talker (same, different), and the interaction between them. ERP models also included fixed main effects of Masker type and Talker (same, different), but also included a main fixed effect of Word Type (color, object), as well as all interactions between these three factors. All models included random by-participant intercepts. All models were implemented in R version 4.2.1 (R Core Team, 2024) and fit using the “lmer” function in the lme4 library (Bates et al. 2015). When significant effects were identified, we conducted post hoc Bonferroni-adjusted pairwise comparisons as follow-up tests using the *emmeans* package (Lenth and Piaskowski, 2025).

## III. Results

### A. Behavior

In both experiments, when the competing masker stream comprised intelligible words, the hit rate was lower when target and masker were the same talker compared to when they were different (Fig. 3, panels A and B: blue lines slant upward to the right). However, when the masker contained unintelligible scrambled words, talker identity had little effect on hit rate (Fig. 3, panels A and B: salmon lines are relatively flat). Relatedly, in both experiments, when the talkers were the same, hit rate was lower for the word masker than the scrambled masker (Fig. 3, panels A and B: salmon points are above blue points on the left side of each panel), but when the talkers differed, the hit rates were more similar for word and scrambled maskers (Fig. 3, panels A and B: salmon and blue points are similar on the right side of each panel).

**Figure 3.**
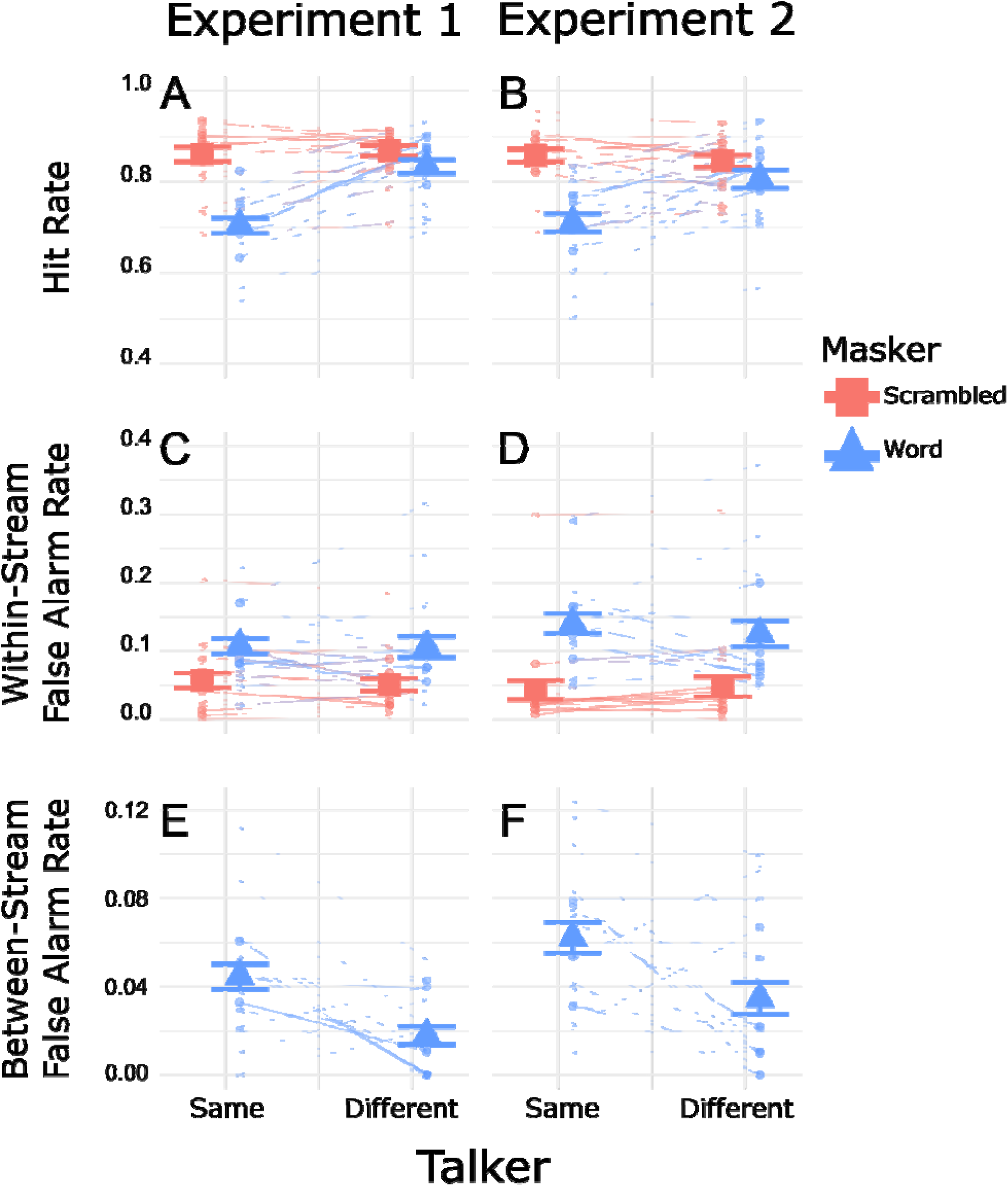
Behavioral results showing individual data (small symbols connected by thin lines) and across-participant average and standard error (large symbols with error bars). Scrambled masker results are in salmon squares, and Word masker results are in blue triangles. **A:** Hit rates for Experiment 1. **B:** Hit rates for Experiment 2. **C:** Within-stream false alarm rates for Experiment 1. **D:** Within-stream false alarm rates for Experiment 2. **E:** Between-stream false alarm rates for Experiment 1. **F:** Between-stream false alarm rates for Experiment 2.

Statistical analyses supported these observations. For Experiment 1, the linear effects model on hit rates revealed significant main effects of Masker (F(1,57) = 150.07, p < 0.0001) and Talker (F(1,57) = 78.06, p < 0.0001), as well as a significant interaction between them (F(1,57) = 61.53, p < 0.0001). To interpret the significant interaction, we ran separate tests to see if there were significant differences between hit rates for word and scrambled maskers for the different masker talker types. When the target and masker talkers were the same, the hit rate was significantly lower for the word masker than the scrambled masker (p < 0.001). When the target and masker talkers differed, the hit rates were also statistically different (p = 0.003), but the effect size was much smaller. In Experiment 2, we also observed significant main effects of Masker (F(1,57) = 66.98, p < 0.0001) and Talker (F(1,57) = 13.26, p < 0.001), as well as a significant interaction (F(1,57) = 22.51, p < 0.0001). In Experiment 2, the hit rate was significantly lower for the word masker than the scrambled masker both when the talkers were the same (p < 0.0001) and when they were different (p = 0.018). Again, the effect of masker type was more drastic when the talker was the same.

Within-stream false alarm rates, i.e., the rate at which participants incorrectly responded to object words within the target stream, were higher for the word masker compared to the scrambled masker (Fig. 3, panels C and D: blue points are higher than salmon points in all cases). In Experiment 1, there was little effect of the masker talker identity for either the word or the scrambled masker (Fig. 3, panel C: both blue and salmon lines are relatively flat). The same was true for Experiment 2: Masker talker identity had little effect, but within-stream false alarm rates were higher for the word masker than for the scrambled masker.

Statistical tests on the within-stream false alarm rate found support for these observations. For Experiment 1, the linear mixed effects model revealed a significant effect of Masker type (F(1,57) = 64.37, p < 0.0001), while the effect of Talker was not significant (F(1,57) = 0.352, p = 0.555), and neither was the interaction between them. In Experiment 2, there was a significant main effect of Masker (F(1,57) = 105.96, p < 0.0001), but not a significant effect of Talker (F(1,57) = 0.318, p = 0.575), nor a significant interaction between them.

In both experiments, false alarms to color words in the masker stream were extremely low (less than 12%; Fig. 3, panels E and F). However, in both experiments these false alarm rates were higher when the target and masker were spoken by the same talker compared to when they differed (Fig. 3, panels E and F: lines slant down to the right in both panels).

Linear-effects models for the between-stream false alarm rate substantiate these observations. In both experiments, there was a significant main effect of Talker (Experiment 1: F(1,19) = 17.56, p < 0.001; Experiment 2: F(1,19) = 12.071, p = 0.0025). This confirms that between-stream false alarm rates were significantly higher for the same talker compared to the different talker condition in both experiments.

### B. Sensory responses: ERP to Masker

In Experiment 1, the word masker had sharp onsets every 300 ms, but the scrambled speech masker did not. The resulting ERPs thus contained strong masker-driven responses for word maskers, but not for scrambled maskers, which complicates interpretation of any differences in target-driven ERPs across the two masker conditions. We ran Experiment 2, with word-modulated scrambled maskers, as a control, as we expected the masker-driven ERPs to be similar for the two masker types, since both had similar, strong rhythmic structure.

The ERPs time-locked to the maskers (Fig. 4, left panels) confirm that in Experiment 1, word maskers evoked strong, rhythmic signals, but scrambled maskers evoked little consistent response. In contrast, in Experiment 2 (Fig. 4, right panels) both word-modulated scrambled maskers and word maskers led to frontocentral EEG responses that were essentially identical.

**Figure 4.**
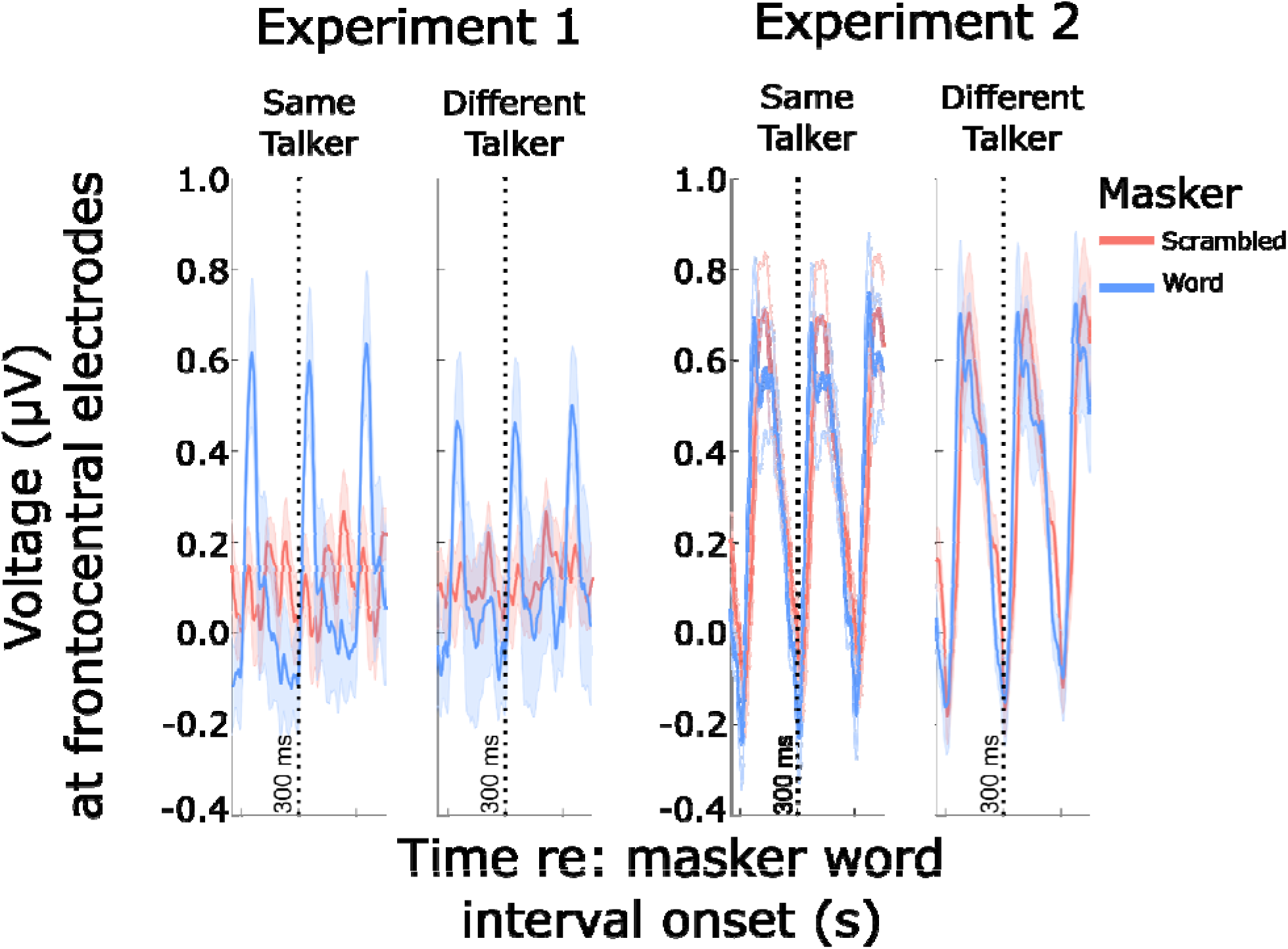
ERPs evoked by masker streams over 900 ms-long epochs over frontocentral electrodes (time locked to word onsets in the word masker). **Left column:** ERPs for Experiment 1. For the word maskers, there are clear rhythmic patterns repeating every 300 ms, corresponding to the onsets of successive words (blue lines in both panels show repeating temporal structure, reflecting word onsets). For the unmodulated scrambled masker, no clear temporal structure is present in the ERPs (red lines in both panels show no strong signal). **Right column:** ERPs for Experiment 2. Both the word masker and the word-modulated scrambled masker show strong, rhythmic ERPs, reflecting the fluctuations in the masker energy over time.

We also considered that ERPs to words with plosive onsets (like “blue”, “green”) are sharper than those to non-plosives (like “red”, “white”). This comparison is shown for ERPs to target words in Supplementary Figure A. Both color and non-color words have a similar percentage of plosive (“bag”, “desk”, “glove”, “pen”, “table”, “toy”) and non-plosive (“hat”, “card”, “chair”, “shoe”, “sock”, “spoon”) onset words. Although we observed that plosive words resulted in sharper, more well-resolved ERPs, the number of plosive and non-plosive words were relatively balanced in color and object words and so did not likely affect the comparisons across Masker and Talker type. There were no significant differences in ERPs to color versus non-color words in the masker.

Only ERPs corresponding to hits and correct rejections were included in our target ERP analysis. Supplementary Fig. B shows ERPs to target color words sorted by hits and misses, as well as ERPs to target object words sorted by false alarm and correct rejections. Note, however, that only hits and false alarms were used in calculating behavioral outcomes. In general, hits elicited stronger P1-N1 and P300 than did false alarms. There were no substantial differences in ERPs corresponding to false alarms versus correct rejections. Additional analyses show that including incorrect answers does not meaningfully alter the results (Supplementary Material C).

### C. Sensory responses: P1-N1 to target stream words

In both experiments, the P1-N1 response was larger (i.e. deeper in the time trace) in response to a target color word than in response to a target object word (Fig. 5). Additionally, P1-N1 responses to target words were larger with a Scrambled masker than a Word masker.

**Figure 5.**
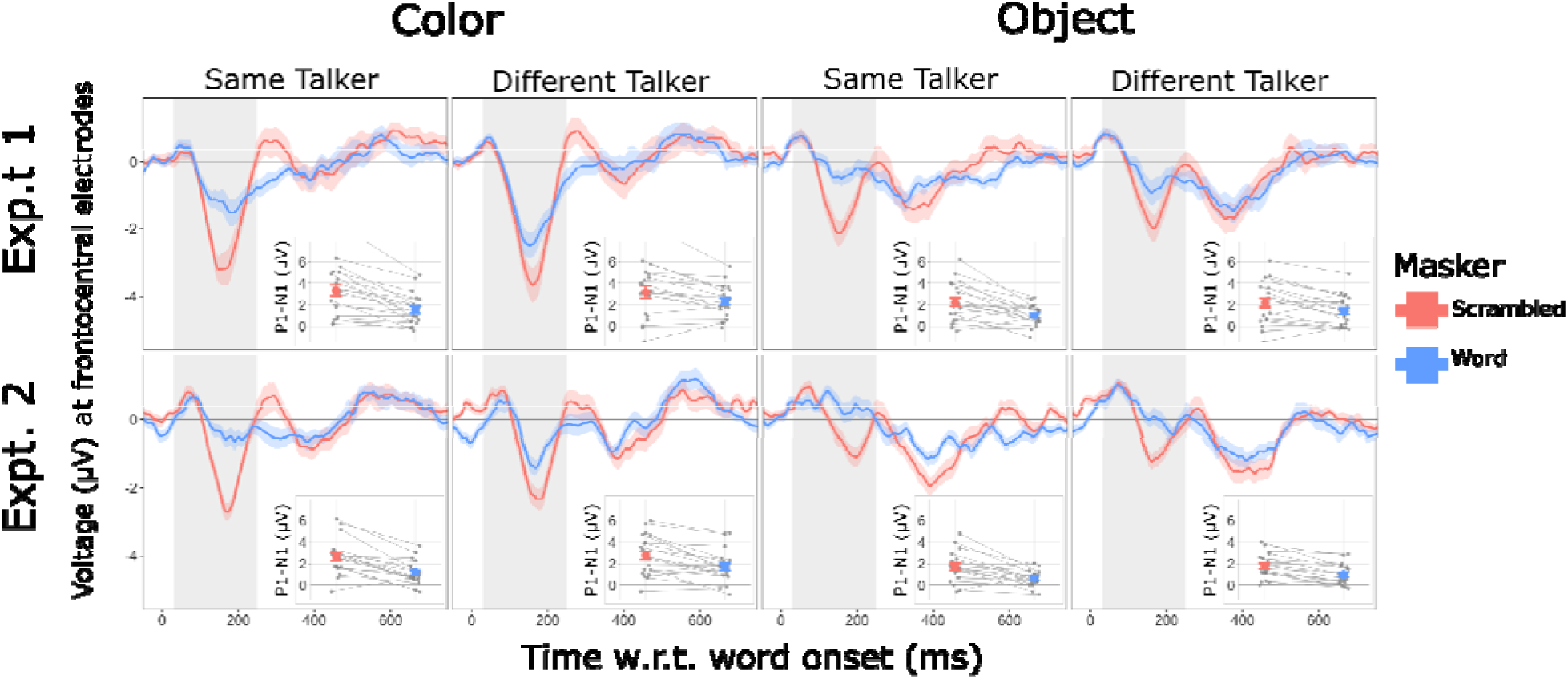
Target-word evoked ERPs over frontocentral electrodes for Experiment 1 (top row) and Experiment 2 (bottom row). Columns show ERPs to target color words (left two columns) and target object words (right two columns) and produced either by the same talker (first and third columns) or different talker (second and fourth columns). ERP time traces are averaged over participants, with bands showing the across-participant standard error of the mean. Inset plots within each panel show P1-N1 magnitudes. Group average and standard error of the mean shown by large symbols. Individual participant data are connected by thin lines.

Statistical analysis supported these observations. In experiment 1, the model on P1-N1 showed significant main effects of Masker (F(1,133) = 64.7, p = 4.235e-13), Talker (F(1,133) = 7.37, p = 0.0075), and Word Type (F(1,133) = 47.28, p = 2.176e-13). The interaction between Masker and Talker was significant (F(1,133) = 5.66, p = 0.019). The interaction between Masker and WordType was marginally significant (F(1,133) = 3.55, p = 0.06). The main effect of Word Type confirms that P1-N1 was significantly larger in response to color words in the target stream than object words. To interpret the significant interaction between Masker and Talker, we tested whether there was a significant difference between P1-N1 for same versus different talkers at each level of masker type. With a Word masker, P1-N1 was significantly higher when the talker was different than when it was the same (p = 0.0004). However, with a scrambled masker, there was not a significant difference (p = 0.813).

The patterns observed in experiment 1 are replicated in experiment 2. The linear mixed effects model on P1-N1 revealed significant main effects of Masker (F(1,133) = 94.8, p = 2.2e-16), Talker (F(1,133) = 8.56, p = 0.0040), and WordType (F(1,133) = 51.30, p = 4.867e-11). The interaction between Masker and Talker was also significant (F(1,133) = 5.24, p = 0.023). The interaction between Masker and WordType was marginally significant (F(1,133) = 3.22, p = 0.075). The main effect of Word Type confirms that P1-N1 was significantly larger in response to color words in the target stream. To interpret the significant interaction between Masker and Talker, we tested whether there was a significant difference between P1-N1 for same versus different talkers at each level of masker type. With a Word masker, P1-N1 was significantly higher when the talker was different than when it was the same (p = 0.0003). However, with a scrambled masker, there was not a significant difference (p = 0.653).

In addition to the linear mixed effects model analysis, we also conducted a post hoc linear correlation analysis between individual behavioral performance and P1-N1 magnitudes, shown in Supplementary Material D. Significant, although relatively weak, positive correlations existed between hit rate and P1-N1 magnitudes in Experiment 1. There was additionally a significant negative relationship between between-stream false alarm rate and P1-N1 in Experiment 2.

### D. Recognition responses: P300 to target stream words

As shown in Figure 6, P300 was larger in response to color words than object words, and for object words was often near zero (i.e. object words did not elicit a P300 response). In contrast to the P1-N1, the P300 was largely unaffected by talker or masker type condition, except for in experiment 1. These patterns were supported by the linear mixed effects model results. The linear mixed effects model on P300 in Experiment 1 showed significant main effects of Masker (F(1,133) = 8.4658, p = 0.0042) and Word Type (F(1,133) = 86.903, p = 3.264e-16). The interaction between Masker and Talker was also significant (F(1,133) = 6.05, p = 0.015). The main effect of Word Type confirms that the P300 elicited by color words was larger than that to object words. To test the interaction between masker and talker, we compared scrambled and words P300 at each level of talker. For a same talker masker, there was a significantly larger P300 to target words with a Scrambled than a Word masker (p = 0.0002). This was not the case for a different talker masker, where P300s for Scrambled and Word maskers were not significantly different (p = 0.751).

**Figure 6.**
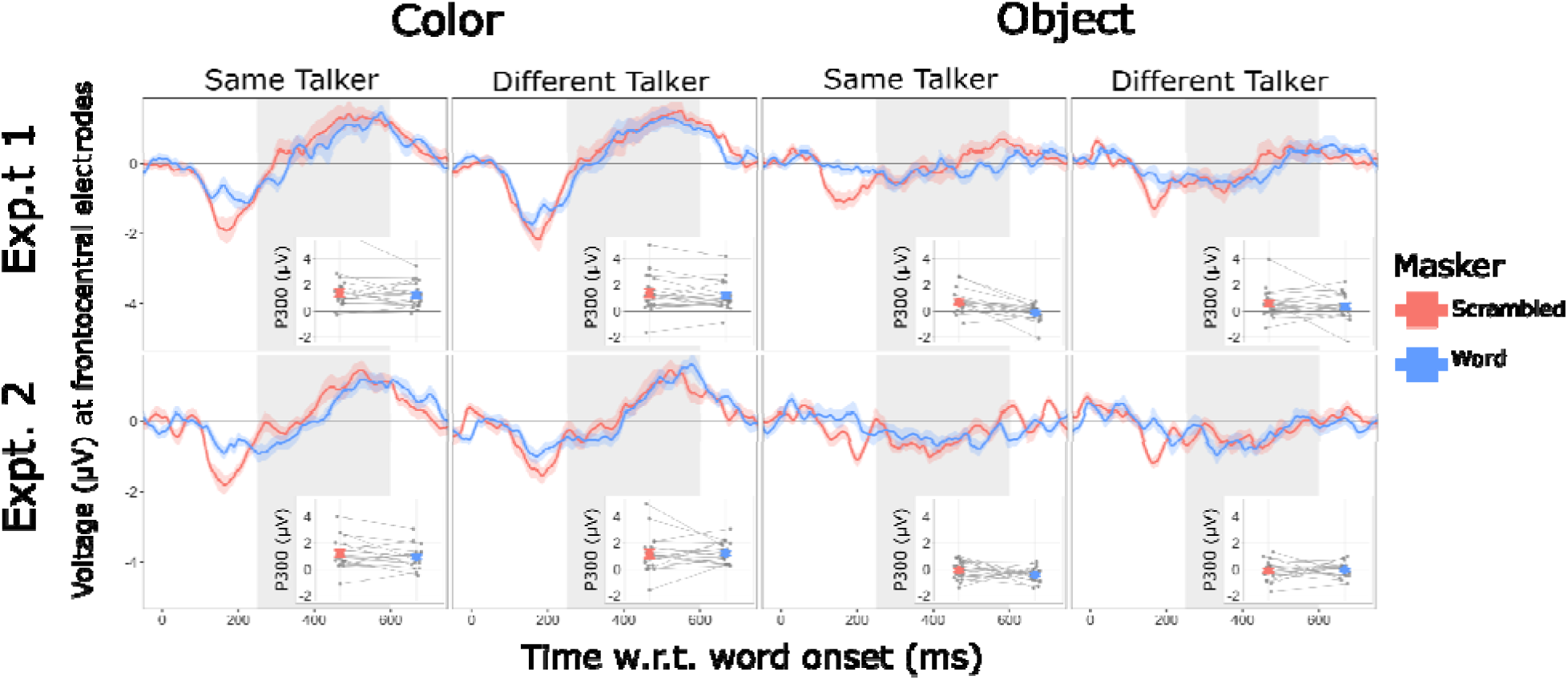
Target-word evoked ERPs over parietooccipital electrodes for Experiment 1 (top row) and Experiment 2 (bottom row). Columns show ERPs to target color words (left two columns) and target object words (right two columns), and produced either by the same talker (first and third columns) or different talker (second and fourth columns). ERP time traces are averaged over participants, with bands showing the across-participant standard error of the mean. Inset plots within each panel show P300 magnitudes. Group average and standard error of the mean shown by large symbols. Individual participant data are connected by thin lines.

For experiment 2, the LMEM revealed only a significant effect of Word Type (F(1,133) = 146.6063, p < 2e-16). This main effect confirms that P300s to color words were larger than those to object words.

As with the sensory P1-N1, we conducted a correlation analysis between individual behavioral performance and P300 magnitude, shown in Supplementary Material D. We found positive correlations whose linear fit was significant between hit rate and P300 in both experiments and for both the Scrambled and Words masker. There were also loosely negative relationships between P300 and within-/between-stream false alarm rates, but none of these were a significant linear fit.

## IV. Discussion

Both experiments in this study tested how masker intelligibility and talker similarity impact selective auditory attention to a sporadically timed target sound presented alongside an ongoing, isochronous masker. Behavioral and neural data converged in showing that selective attention to the target was most impaired when the masker was hardest to ignore: when it was intelligible and spoken by the same talker. This is evidenced by lower hit rates, higher within-and between-stream false alarm rates, and reduced sensory P1-N1 responses in the EEG. In contrast, target recognition P300 responses were primarily modulated by target word category (color vs. object) and showed little sensitivity to masker type or talker, suggesting that masker-related interference predominantly affected early stages of sensory-perceptual processing rather than later recognition stages.

All of these effects were observed under conditions designed to minimize energetic masking and to match key spectrotemporal properties between Word and Scrambled maskers. Although scrambling disturbed phoneme and word-level frequency information, local (within 250 ms) spectrotemporal information and pitch were largely preserved. However, these manipulations do not eliminate all possible acoustic differences, and thus cannot fully dissociate effects of intelligibility from residual acoustic cues. Still, within these constraints, our results are consistent with a contribution of intelligibility and talker similarity to masking beyond what would be expected from reduced acoustic differences alone. More broadly, the results align with prior work showing that intelligible competing speech can disrupt perception and cortical responses even when low-level acoustic differences are minimized. Critically, the dissociation between P1–N1 and P300 responses suggests that intelligibility-related masking primarily limits the likelihood that target sounds are successfully selected for further processing, rather than degrading recognition once selection has occurred.

### A. Intelligible speech more readily interrupted selective attention than scrambled speech, despite spectrotemporal similarity

Intelligible masking speech interrupted target color word detection much more readily than scrambled speech. Participants showed higher hit rates and lower within-stream false alarm rates with a Scrambled than a Word masker, consistent with many previous findings that intelligible maskers impair performance more strongly than unintelligible maskers. However, previous work varied in how differences in intelligibility between interferers were imposed. Examples include comparing speech and noise directly (Brungart and Simpson, 2002), varying the number of talkers in an N-talker background (Kurthen et al., 2021; Van Engen et al., 2014), presenting maskers in different languages (Van Engen and Bradlow, 2007), filtering the interferer (Pollack, 2005; Rennies et al., 2019), or noise vocoding (Scott et al., 2006). Many of these manipulations do not address the confound that intelligible vs. unintelligible speech maskers might also vary in the degree to which they energetically mask the target sound.

Here, target and masker streams were filtered into interleaved, non-overlapping frequency bands and resummed, dramatically reducing energetic overlap. Additionally, scrambling procedures equated spectrotemporal density across maskers. Finally, Experiment 2 imposed identical amplitude modulation on both masker types, reducing differences in envelope cues over time. These results suggest that intelligibility contributes to masking beyond what can be explained by energetic and spectrotemporal factors, which were substantially reduced but not entirely eliminated in the present design. Studies that attempt to control for spectrotemporal differences between different masking sounds (Strelnikov et al., 2011; Summers and Roberts, 2020), like the present findings, report persistent intelligibility effects even when acoustic overlap is minimized. Similar arguments appear in work showing that competing intelligible speech remains a potent masker even when its long-term spectrum is matched to speech-shaped noise and energetic masking is reduced (Brungart et al., 2006; Miller and Licklider, 1950; Summers and Roberts, 2020).

### B. The effect of masker intelligibility was more pronounced when target and masker were spoken by the same talker

Behavioral effects of masker intelligibility were exaggerated when target and masker were spoken by the same talker. This might tell us about the ease of the different talker condition: when the masker was spoken by a different talker, it was trivial to hear out target color words (achieve high hit rate and low within-stream false alarm rate) regardless of whether the masker was intelligible or not. Conversely, in the same talker case, participants showed a large difference between Word and Scrambled maskers, implying that a same-talker Word masker provided a substantial disruption to attention to the target. This result agrees with work showing that similarity between intelligible competing sounds, particularly shared vocal characteristics like pitch and vocal tract length, enhances masking (Brungart et al., 2001, 2006; Darwin, 2008). Additionally, the between-stream false alarm rate data show that participants were more likely to mistakenly respond to masker color words with the same talker masker. When talker properties are shared, stream segregation is weakened (Best et al., 2007; Richardson et al., 2025; Shinn-Cunningham et al., 2017), increasing the likelihood that masker events are perceived as belonging to the target.

### C. Sensory P1-N1 responses were modulated by masker intelligibility, talker differences, and word type

Cortically, P1-N1 responses to targets were smaller with a Word masker than with a Scrambled masker. With a Word masker, there was an effect of talker: P1-N1 was significantly higher when the talker was different. This was not the case when the masker was Scrambled, where there was not a significant difference between same and different talker. In the present context, a smaller P1-N1 indicates that same-talker intelligible maskers prevented auditory attention from strongly modulating the target response. The P1–N1 modulation may reflect enhanced sensory gain for target-relevant categories, consistent with attention-driven modulation of early auditory cortical responses. It is well known that attention modulates early cortical responses in favor of attended sound events, and that this process involves top-down modulation from the prefrontal cortex. This is evidenced by studies from the neuronal level in animals (Hockley and Malmierca, 2024; Macedo-Lima et al., 2024; McGinley et al., 2015; Niwa et al., 2012, 2015; Winkowski et al., 2018) to non-invasive neuroimaging measures in humans (Alho et al., 2014; Choi et al., 2014; Hillyard et al., 1973; Lee et al., 2013; Woldorff et al., 1993). Here, the difference between P1-N1 to color words across masker conditions elucidates the attentional mechanism at play. When attention to the target was more difficult (Word masker, same-talker masker), cortical sensory responses were smaller. Additionally, when the sound event was to-be-attended (i.e., a color word), it drove a larger sensory response than a to-be-ignored event (i.e., an object word).

This pattern is also supported by our linear correlation analysis (Supplementary Material D). A positive correlation between hit rate and P1-N1 implies that participants with a greater degree of attention-drive modulation of early sensory responses perform better on the task. This is consistent with previous reports that both P1-N1 and neural activity in the superior temporal gyrus correlates with task performance (Choi et al., 2014; Zhang et al., 2021). However, this relationship may also reflect individual differences in task engagement, whereby participants who are more engaged with the task exhibit less movement and consequently yield more robust ERP responses.

Given that release from energetic masking strongly facilitates segregation (Brungart et al., 2006; Oxenham and Simonson, 2009), energetic masking is always a concern when acoustic manipulation of the target or masker is involved. However, our scrambling procedure carefully controlled for differences between Word and Scrambled maskers such that they masked the target to a similar degree. Tantalizingly, we still observed effects of attention with these controls in place, arguing against purely sensory-driven explanations for these results; Intelligibility and talker identity shaped performance and sensory responses beyond acoustic effects. This reinforces the idea that high-level attributes like speech intelligibility and talker identity can dominate perception and sensory responses even when low-level cues favor segregation. Intelligibility exerts its influence at a sensory processing stage, consistent with ideas that linguistic interference, semantic competition, and attention capture by meaningful speech disrupt selective attention (Bregman, 1990, 2003; Mattys et al., 2012; Shinn-Cunningham, 2008).

The persistence of P1-N1 effects in Experiment 2, when a 3.33 Hz modulation drove a strong onset response (Fig. 4), indicates that P1-N1 differences were not confounded by ongoing periodicity (or lack thereof) in the Scrambled masker.

### D. Color words drove strong P300 target recognition responses regardless of masker type

Color words, when successfully recognized, drove strong P300 responses compared to object words. There was a relatively small effect of masker type on P300 to object words in the same talker condition (Exp. 1), but this is not likely meaningful - P300 responses to object words were near zero in general. This difference is consistent with extensive work showing that P300 is elicited by relatively rare, attended, task-relevant sound events (Başar-Eroglu et al., 2001; Collard et al., 2008; Duncan-Johnson and Donchin, 1982; Picton, 1992; Polich, 2007). For this task, participants were required to categorize each word in the target stream (color or object), and the P300 signals elicited by color words represent successful recognition of word type. Additionally, our correlation analysis showed that participants who performed better at the task had larger P300s than the worse performers. However, within each trial, color words were presented with equal frequency to object words, likely meaning that these P300s were smaller than they would be for a lower color word probability (Kurt et al., 2025; Polich, 2007).

Because only ERPs to correctly identified color words were included in the P300 analysis, our P300 effects likely reflect the neural signature of successful target recognition. Thus, the absence of masker effects on P300 does not imply that recognition itself is unaffected; rather, intelligible and same-talker maskers likely reduce the probability that targets reach the recognition stage at all. An open question is whether the presence of a masking sound in general affects P300 amplitudes, as it does indeed lead to lower likelihood of target sound recognition.

The P300 results directly contrast with the sensory P1-N1 results, which showed strong effects of masker type, word type, and talker identity. The dissociation observed here, showing strong intelligibility and talker effects on P1–N1 but not P300, supports the idea that masker intelligibility disrupts early sensory selection. When selection succeeds, however, recognition proceeds robustly and independently of masker characteristics.

The present neural results confirm that intelligible, same talker masking sounds disrupt selective attention at early sensory stages, suggesting that the ability to ignore distracting speech depends strongly on sensory-perceptual parsing rather than solely on later cognitive control.

## V. Conclusion

In this study, intelligible speech maskers, particularly when spoken by the same talker as the target, substantially disrupted selective auditory attention even when energetic masking and spectrotemporal properties of the masker were tightly controlled. A same talker, intelligible masker resulted in poor behavioral performance and reductions in early sensory (P1-N1) responses. In contrast, P300 responses indexing successful target recognition were largely insensitive to masker intelligibility and talker differences, suggesting an attentional filtering mechanism at the sensory level rather than the level of recognition. Taken together, these findings reaffirm that masking by intelligible speech primarily arises from failures of early sensory perceptual filtering.

## Acknowledgements

The authors gratefully acknowledge Emaya Anand and Naomi Jesionowski for their help in data collection and behavioral data analysis, and Misha Psenicka for their helpful comments during data analysis and preparation of this manuscript. This work was supported by grants from the Office of Naval Research N00014-23-1-2065, National Institute on Deafness and Other Communication Disorders R01 DC019126 to BGS and MJR, National Institute on Deafness and Other Communication Disorders R01 DC022699 to CAB and BGS, and National Institute on Deafness and Other Communication Disorders F31 DC022481 to BNR.

## VI. Author Declarations

### B. Ethics Approval

All participants in this study gave written informed consent prior to participation. All testing was administered according to the guidelines of and with approval from the Institutional Review Board of Carnegie Mellon University.

### C. Data Availability

The data and analysis supporting the conclusions of this manuscript are available upon request from the authors.

