## Supplementary Material for "Intelligible distracting speech disrupts early auditory attention"

### Supplementary Materials

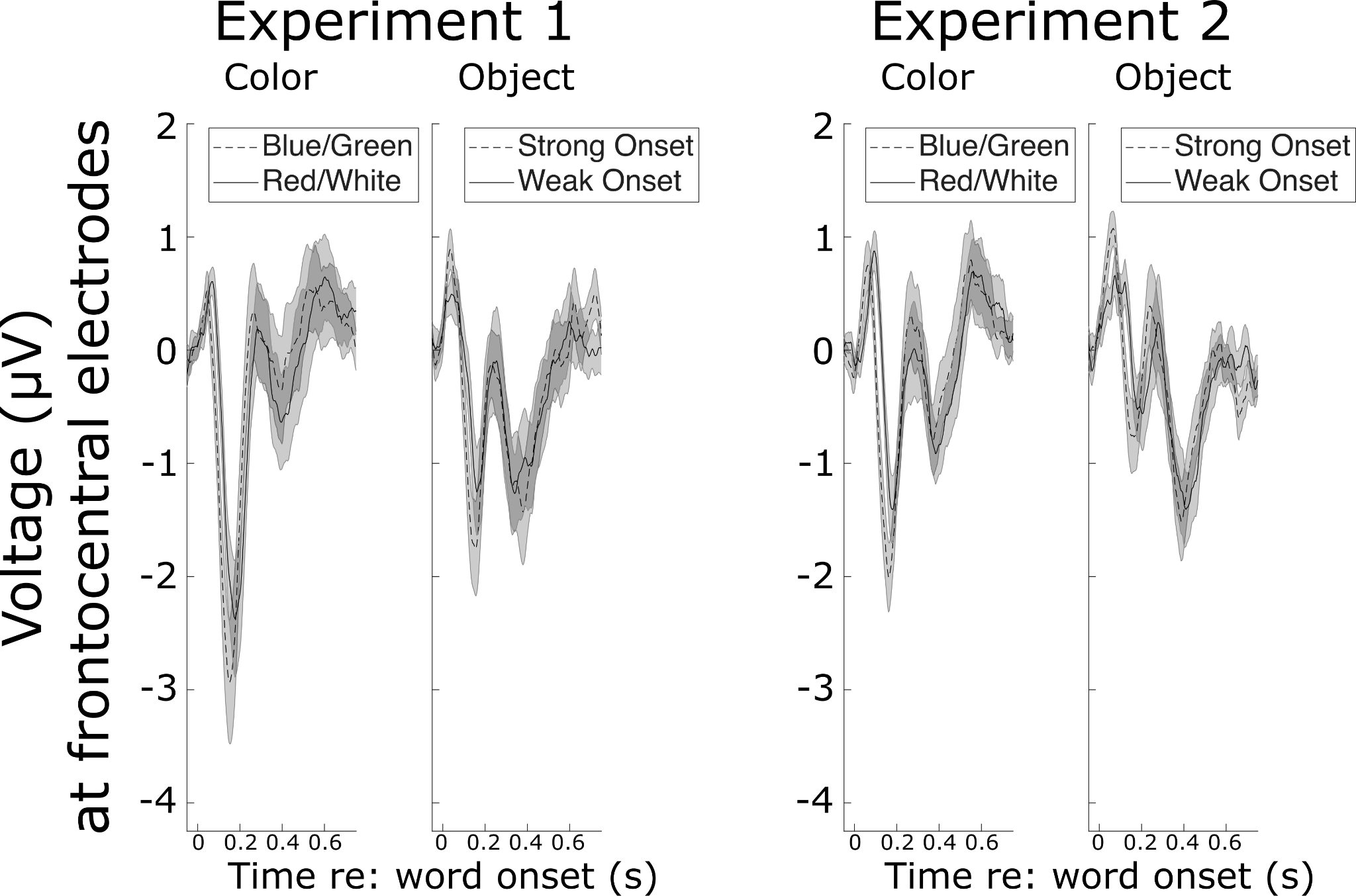

**Supplementary Figure A**. Frontocentral ERPs to plosive vs. non-plosive words. Left panel in each Experiment: ERP to target color words sorted by plosive (blue, green) versus non-plosive (red, white) words. Right panel in each Experiment: ERP to target object words sorted by plosive words (“bag”, “desk”, “glove”, “pen”, “table”, “toy”) verus non-plosive words (“hat”, “card”, “chair”, “shoe”, “sock”, “spoon”).

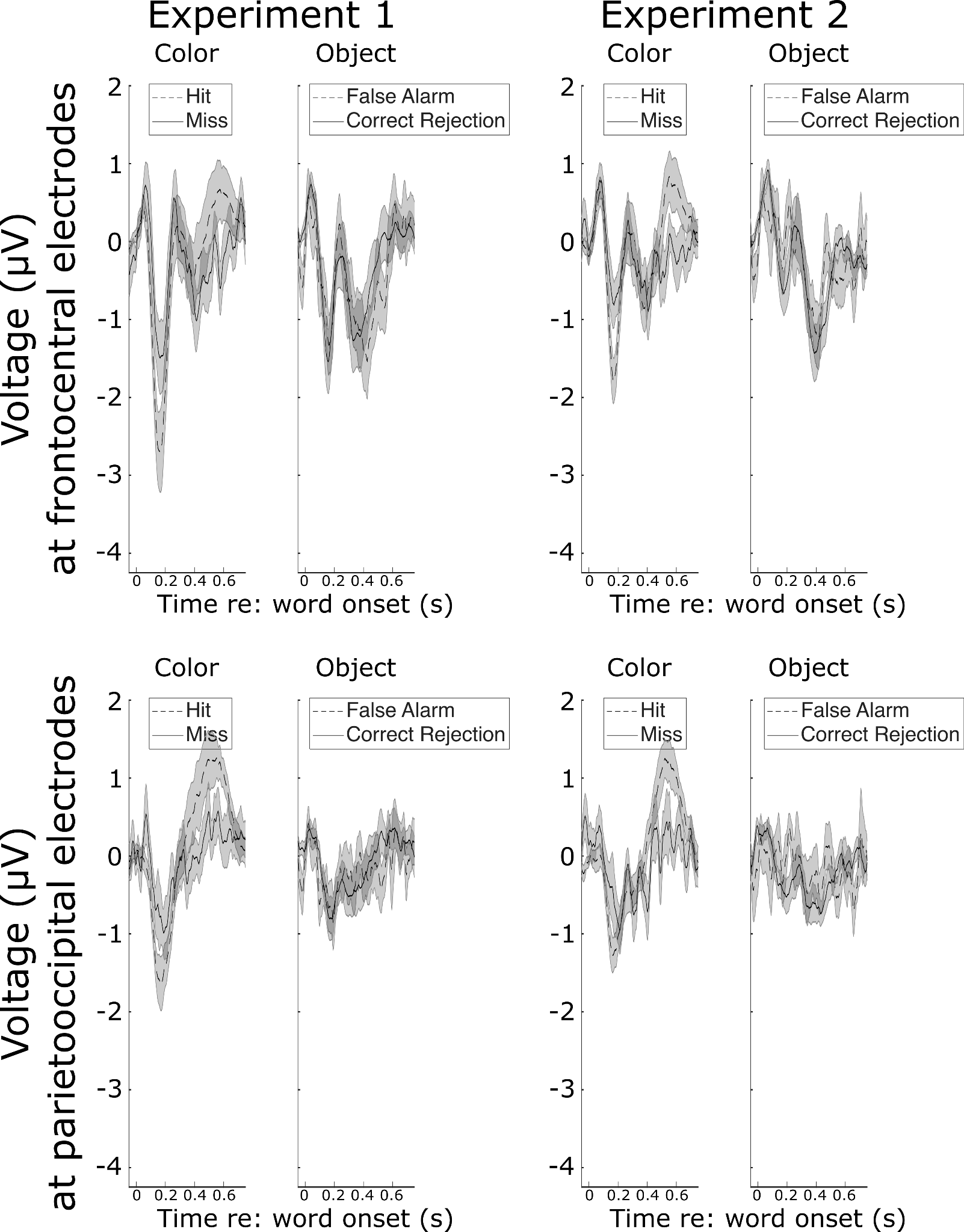

**Supplementary Figure B.** ERPs to target color words sorted by response type. Top Row: Frontocentral ERPs to target color words sorted by hits (responded) versus misses (did not respond) in the left panel for each Experiment. Frontocentral ERPs to target object words sorted by false alarms (responded) versus correct rejections (did not respond) in the right panel for each experiment. Bottom Row: Parietooccipital ERPs to target color words sorted by hits (responded) versus misses (did not respond) in the left panel for each Experiment. Parietooccipital ERPs to target object words sorted by false alarms (responded) versus correct rejections (did not respond) in the right panel for each experiment.

##### Supplementary Material C

In the main manuscript, we report ERP results which only include responses to words that correspond to hits and correct rejections (that is, misses and false alarms are not included). Below, we describe the statistical analysis conducted which includes misses and false alarms to test whether this meaningfully affected the results. When ERPs corresponding to incorrect responses are included, the results remain largely unchanged. There are two exceptions: the significant interaction between Masker and Talker for P1-N1 magnitudes in experiment 2 is not significant when incorrect responses are included, and the P300 data in experiment 1 do not show a significant interaction between Masker and Talker. This agrees with the trend in data plotted in Supplementary Fig. B, where including misses and false alarms introduces noise.

#### P1-N1

In experiment 1, the model on P1-N1 showed significant main effects of Masker (F(1,133) = 63.6, p = 6.12e-13) and Word Type (F(1,133) = 35.38, p = 1.16e-8). The interaction between Masker and Talker was significant (F(1,133) = 5.66, p = 0.02). The main effect of Word Type confirms that P1-N1 was significantly larger in response to color words in the target stream than object words. To interpret the significant interaction between Masker and Talker, we tested whether there was a significant difference between P1-N1 for same versus different talkers at each level of masker type. With a Word masker, P1-N1 was significantly higher when the talker was different than when it was the same (p = 0.0062). However, with a scrambled masker, there was not a significant difference (p = 0.611).

In experiment 2, the linear mixed effects model on P1-N1 revealed significant main effects of Masker (F(1,133) = 77.17, p = 6.86e-15), Talker (F(1,133) = 4.91, p = 0.02), and WordType (F(1,133) = 39.9, p = 3.69e-9). The main effect of Word Type confirms that P1-N1 was significantly larger in response to color words in the target stream. The main effect of Masker confirms that P1-N1 was larger when the masker was Scrambled than Words. Finally, the main effect of talker shows that P1-N1 was larger when the talker was different than when it was the same talker.

#### P300

The linear mixed effects model on P300 in Experiment 1 showed significant main effects of Masker (F(1,133) = 10.57, p = 0.0014) and Word Type (F(1,133) = 71.12, p = 4.91e-14). The main effect of Word Type confirms that the P300 elicited by color words was larger than that to object words. The main effect of Masker confirms that the P300 elicited by color words was significantly larger with a Scrambled masker than a Word masker.

For experiment 2, the LMEM revealed only a significant effect of Word Type (F(1,133) = 121.79, p < 2e-16). This main effect confirms that P300s to color words were larger than those to object words.

##### Supplementary Material D

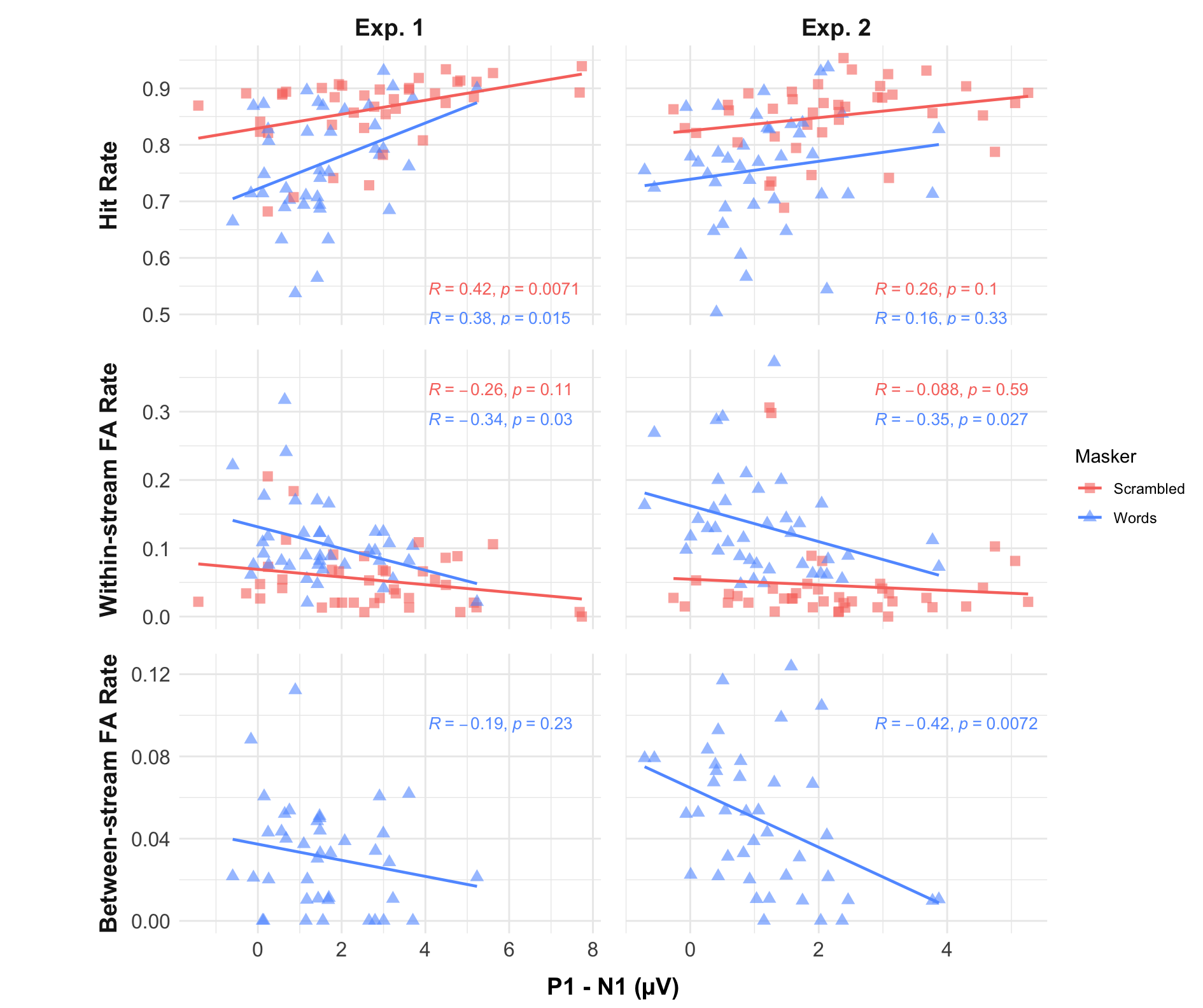

**Supplementary Figure C.** Correlation between behavioral performance and P1-N1 magnitudes. In each panel, each data point shows an individual participant when the masker was Scrambled (salmon squares) or Words (blue triangles). Correlation coefficient R and p-value are listed in the corresponding color in each panel. Top Row: Hit rate vs. P1-N1 for Experiment 1 (Left) and Experiment 2 (Right). Middle Row: Within-stream false alarm rate vs. P1-N1 for Experiment 1 (Left) and Experiment 2 (Right). Bottom Row: Between-stream false alarm rate vs. P1-N1 for Experiment 1 (Left) and Experiment 2 (Right).

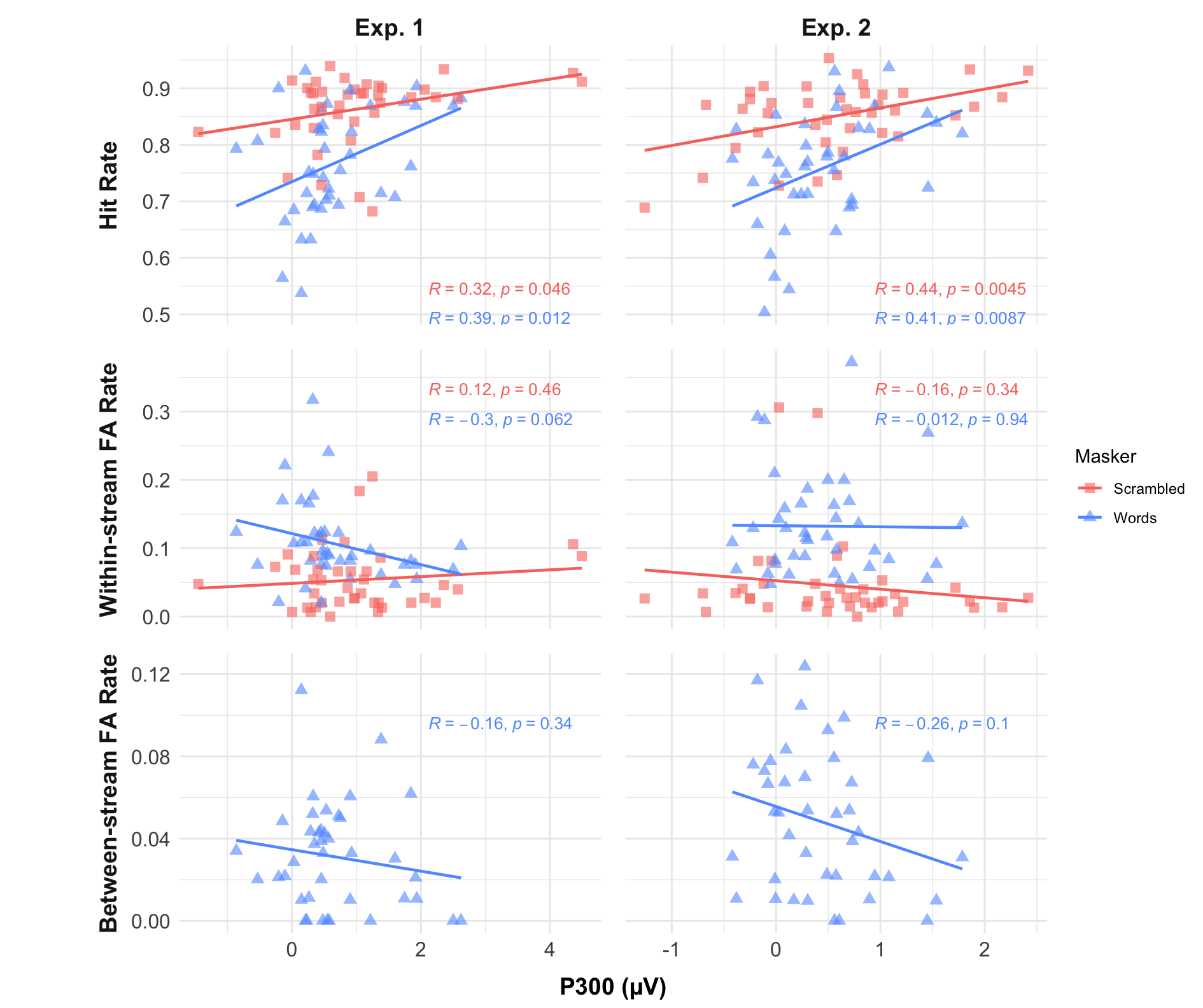

**Supplementary Figure D.** Correlation between behavioral performance and P300 magnitudes. In each panel, each data point shows an individual participant when the masker was Scrambled (salmon squares) or Words (blue triangles). Correlation coefficient R and p-value are listed in the corresponding color in each panel. Top Row: Hit rate vs. P300 for Experiment 1 (Left) and Experiment 2 (Right). Middle Row: Within-stream false alarm rate vs. P300 for Experiment 1 (Left) and Experiment 2 (Right). Bottom Row: Between-stream false alarm rate vs. P300 for Experiment 1 (Left) and Experiment 2 (Right).
